# Characterisation of vaginal *Lactobacillus* isolates from South African women towards the development of a biotherapeutic to optimise the vaginal microbiome

**DOI:** 10.64898/2026.03.26.714511

**Authors:** Jenna Wilson, Aleya Sarah Amir Hamzah, Christine Jordan, Joshua A Hayward, Brian Kullin, Monalisa T Manhanzva, David Tyssen, Celia Mehou-Loko, Andrea G Abrahams, Nina Radzey, Rushil Harryparsad, Bahiah Meyer, Anna C Hearps, Mark Ziemann, Hilton Humphries, Pamela Mkhize, Linda-Gail Bekker, Jo-Ann S Passmore, Heather B Jaspan, Anna E Sheppard, Gilda Tachedjian, Lindi Masson

## Abstract

HIV remains among the world’s most serious healthcare challenges, with adolescent girls and young women in sub-Saharan Africa at particularly high risk of infection. Bacterial vaginosis (BV) is a key risk factor for HIV acquisition, however current treatment strategies are limited. Optimal vaginal *Lactobacillus* spp. protect against BV and HIV, largely through immunoregulatory and antimicrobial activities mediated in part by lactic acid. Towards the development of a *Lactobacillus*-containing live biotherapeutic for African women, we sampled 181 vaginal *Lactobacillus* isolates from 25 BV-negative South African women. Fifty isolates were selected for evaluation of inflammatory responses using vaginal epithelial cells, D- and L-lactate and lactic acid production and culture acidification. Aside from a single *Lactobacillus salivarius* strain, *L. crispatus* isolates acidified the culture media the most and produced the most D- and L-lactic acid. Inflammatory cytokine responses to *Lactobacillus* strains were variable, with *L. crispatus* eliciting the lowest levels of cytokine production. When all properties were evaluated collectively, *L. crispatus* strains exhibited the most desirable biotherapeutic characteristics. Whole genome sequence analysis of ten *L. crispatus* isolates showed that the majority were more closely related to one another than to isolates from other geographical regions. This supports the need for live biotherapeutics to be tailored for the population of intended use. No antimicrobial resistance elements were detected, while putative bacteriocins and intact prophage sequences were identified in all isolates. *L. crispatus* isolates displayed characteristics essential for optimal live biotherapeutic performance, however additional analysis is required to determine the functionality of identified putative prophages.

**Importance:** HIV is a leading cause of morbidity and mortality in sub-Saharan Africa, where adolescent girls and young women are three times more likely to acquire HIV than their male counterparts. A key risk factor for HIV is bacterial vaginosis (BV), a condition characterised by the loss of beneficial *Lactobacillus* species and increased abundance of non-optimal, inflammatory bacteria. Although BV affects approximately 25% of women in sub-Saharan Africa, effective therapeutics are lacking. Live biotherapeutics containing optimal *Lactobacillus* spp. represent a promising strategy to improve BV treatment outcomes and reduce HIV infection risk. We isolated 181 vaginal *Lactobacillus* spp. from 25 BV-negative South African women and characterized 50 selected isolates. This led to the identification of live biotherapeutic candidates for African women with distinct genomes compared to isolates from other geographical regions. This study contributes to current knowledge of the characteristics that should be considered when screening novel isolates for this purpose.

## Introduction

Sub-Saharan Africa (SSA) is the epicentre of the HIV pandemic, where more than 60% of people living with HIV reside (1). In this region, 77% of new infections among people aged 15–24 years are among adolescent girls and young women (1). A key risk factor for HIV infection in SSA women is an imbalance in the vaginal microbiota known as bacterial vaginosis (BV) (2, 3). BV occurs when the relative abundance of optimal *Lactobacillus* spp. decreases with an increase in non-optimal facultative and strict anaerobic bacteria in the female genital tract (FGT) (4). This highly prevalent condition, that affects 42% of 15–24-year-olds and 41% of 25–49-year-olds in South Africa (5), is associated with FGT inflammation and increases risk of sexual HIV acquisition up to 4.4-fold (2, 6–8). In contrast, women who are colonised with optimal *Lactobacillus* spp. have low levels of FGT inflammation and reduced risk of acquiring HIV, other sexually transmitted infections (STIs), and BV (2, 8–11). *L. crispatus*, in particular, has been associated with optimal sexual and reproductive health outcomes, and a previous study in South Africa demonstrated no incident HIV infections among a small number of women with *L. crispatus*-dominant microbiota (2).

Despite the significant role of BV in increasing HIV acquisition risk, and the opportunity to avert HIV infections by reducing BV prevalence in women most at risk of HIV, effective treatment strategies are lacking and the current standard of care, antibiotic treatment, is associated with high BV recurrence rates (12). Although preexposure prophylaxis (PrEP) is a very promising approach to reduce HIV infection rates in key populations, the efficacy of some forms of topical PrEP is reduced in women with BV (13) and multiple studies have demonstrated poor adherence to oral PrEP, particularly in young SSA women at highest risk of HIV (14). The recent approval of the highly efficacious long-acting cabotegravir injectable for HIV prevention in South Africa will likely go a long way to address the limitations of other delivery methods if accessible and acceptable in this population (15). However, broad coverage requires choice, and accordingly the HIV prevention field continues to work towards the development of novel prevention strategies (16).

An adjunctive strategy for BV treatment that may also reduce HIV infection risk is live biotherapeutic administration to recolonize the FGT with optimal *Lactobacillus* spp. Recently a multi-strain *L. crispatus* synbiotic vaginal tablet (VS-01) was found to significantly reduce the relative abundance of non-optimal vaginal microbes and the concentrations of the inflammatory marker, interleukin (IL)-1α, in a randomised, placebo-controlled clinical trial in the United States (17). Similarly a Phase 2b trial conducted in the United States evaluating the impact of a vaginally-derived *L. crispatus* live biotherapeutic product (CTV-05 or LACTIN-V) on BV recurrence showed that BV recurrence was significantly less common among women who received LACTIN-V following antibiotic treatment than in the placebo arm (30% versus 45%, respectively) (18). Another clinical trial conducted in Estonia evaluated the efficacy of two *L. crispatus* strains for BV treatment and found a significant reduction in BV Nugent score as well as clinical signs and symptoms (19). Although the FGT live biotherapeutics field is growing rapidly, there are still no approved biotherapeutics that contain *Lactobacillus* spp. isolated from the FGTs of healthy African women. Previous studies have found that substantial variation in the composition of the vaginal microbiota exists by ethnicity and geographical region (20). *L. crispatus* dominance in African women is relatively uncommon compared to Caucasian and Hispanic women, and large proportions of African women have non-optimal microbiota or sub-optimal *L. iners*-dominant microbiota (8, 21, 22). Furthermore, whole genome sequence (WGS) analysis of *L. crispatus* isolates from various regions of the world demonstrated evolutionary diversity between geographic regions, suggesting that population-specific adaptations may play a crucial role in successful colonisation of the vagina (23). Similarly, the genome sizes and GC content of vaginal lactobacilli have been found to differ from those of the gastrointestinal tract and other sources (24). This suggests that genetic and environmental factors impact bacteria that colonise the vagina and it is thus possible that for colonisation to be effective and sustained, live biotherapeutics should be developed utilising vaginal bacterial strains from women residing in the region of intended use.

In this study we evaluated biotherapeutic-relevant properties of novel vaginal *Lactobacillus* isolates from South African women towards the development of a live biotherapeutic to optimize the vaginal microbiome in African women and potentially reduce HIV infection risk in this population.

## Results

### Lactobacillus isolation

In total, 205 bacteria were isolated from the cervicovaginal secretions of 25 women, including 181 *Lactobacillus* species (72 *L. crispatus,* 30 *L. reuteri,* 24 *L. jensenii,* 23 *L. gasseri,* 22 *L. vaginalis,* 7 *L. johnsonii,* 2 *L. salivarius,* 1 *L. coleohominis)* and other anaerobic bacteria (*Staphylococcus epidermidis, Atopobium vaginae, Streptococcus anginosus, Cutibacterium avidum*). Although *L. iners* is prevalent in this population, this species is considered sub-optimal (4) and was not isolated as additional media supplementation is required. The majority of lactobacilli isolated were *L. crispatus*, followed by *L. reuteri* and *L. jensenii*. From these isolates, 50 lactobacilli from 18 women (aged 16-32 years) were selected based on Gram stain and colony morphologies to include morphologically distinct isolates when selecting more than one of the same species from the same woman (**Table 1**). Due to the high prevalence of STIs (*Chlamydia trachomatis, Trichomonas vaginalis* and *Neisseria gonorrhoeae*) in the parent cohort (53%), four of the women from whom nine *Lactobacillus* isolates were obtained had a STI and these isolates were analysed separately to evaluate whether STI status impacted the characteristics of the isolates.

**Table 1.**
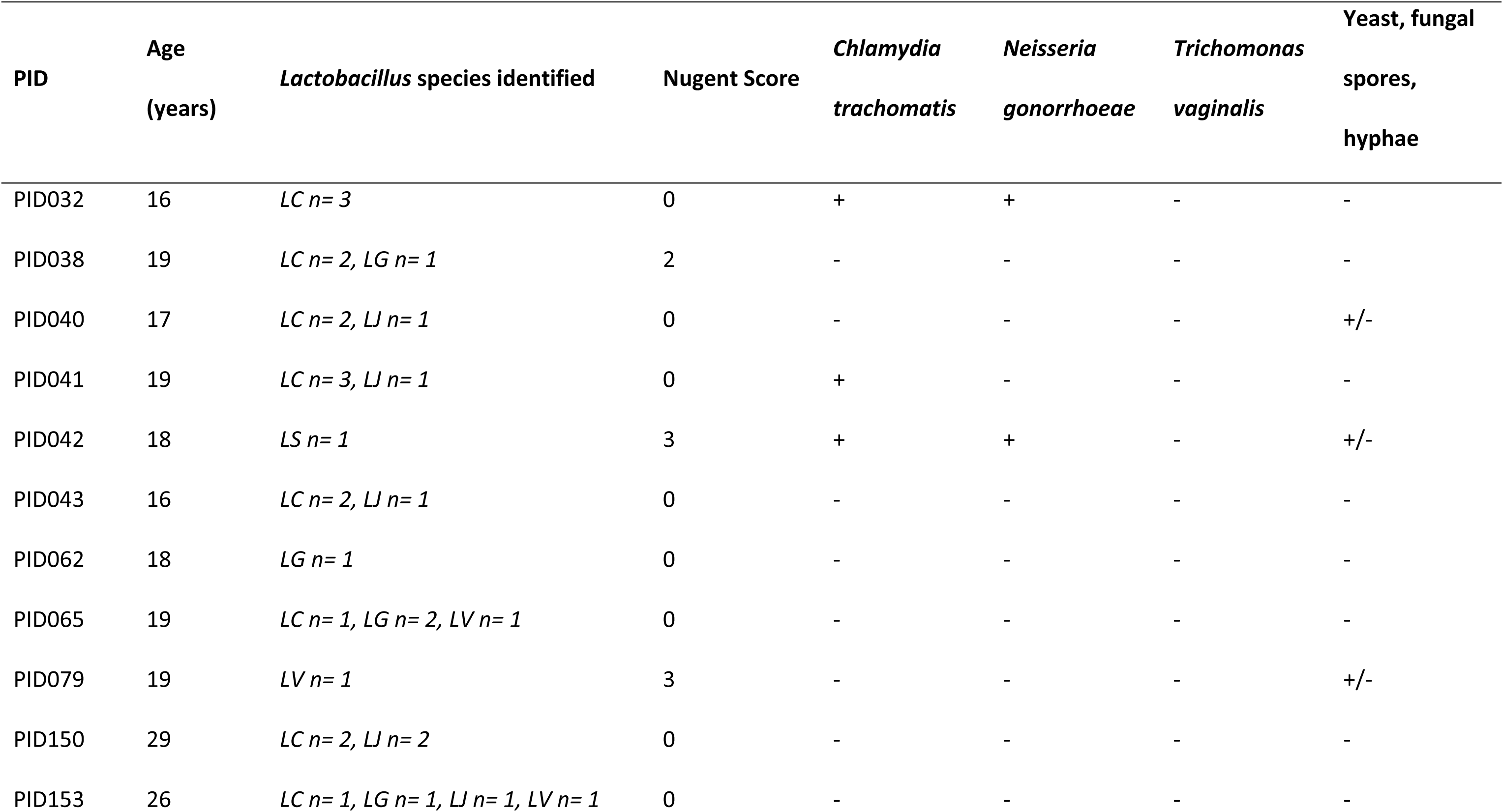
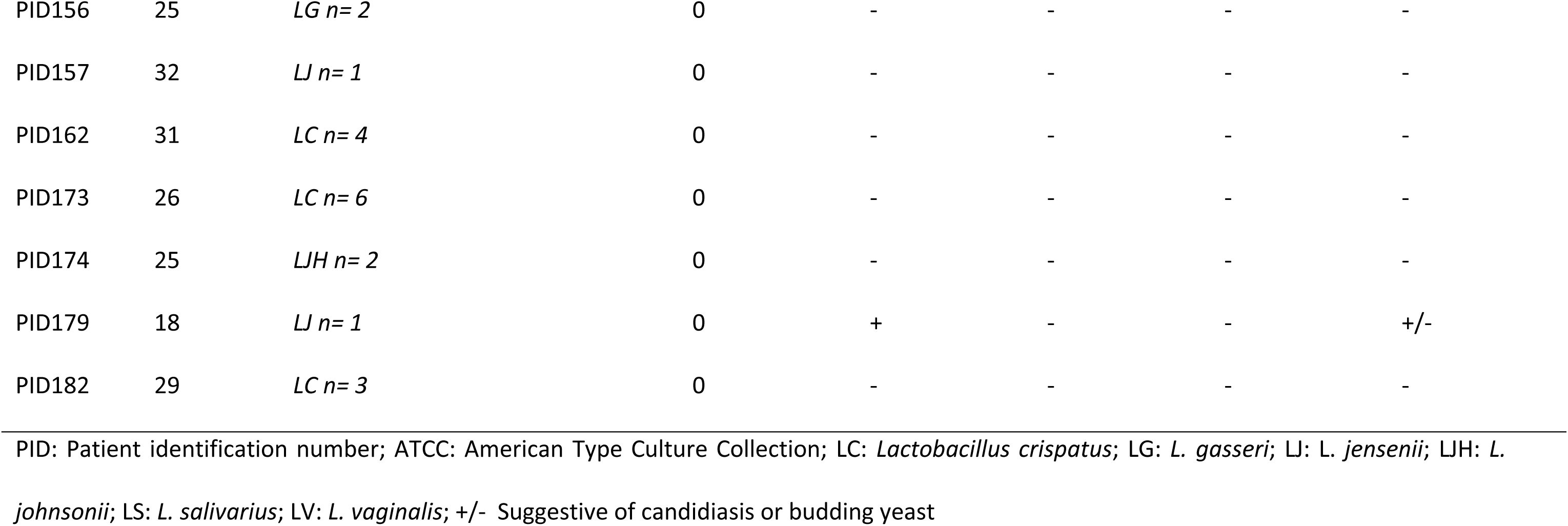
Clinical characteristics of study participants from whom lactobacilli were isolated.

| PID | Age<br>(years) | *Lactobacillus* species identified | Nugent Score | *Chlamydia*<br>*trachomatis* | *Neisseria*<br>*gonorrhoeae* | *Trichomonas*<br>*vaginalis* | Yeast, fungal<br>spores,<br>hyphae |
| --- | --- | --- | --- | --- | --- | --- | --- |
| PID032 | 16 | LC n= 3 | 0 | + | + | - | - |
| PID038 | 19 | LC n= 2, LG n= 1 | 2 | - | - | - | - |
| PID040 | 17 | LC n= 2, LJ n= 1 | 0 | - | - | - | +/- |
| PID041 | 19 | LC n= 3, LJ n= 1 | 0 | + | - | - | - |
| PID042 | 18 | LS n= 1 | 3 | + | + | - | +/- |
| PID043 | 16 | LC n= 2, LJ n= 1 | 0 | - | - | - | - |
| PID062 | 18 | LG n= 1 | 0 | - | - | - | - |
| PID065 | 19 | LC n= 1, LG n= 2, LV n= 1 | 0 | - | - | - | - |
| PID079 | 19 | LV n= 1 | 3 | - | - | - | +/- |
| PID150 | 29 | LC n= 2, LJ n= 2 | 0 | - | - | - | - |
| PID153 | 26 | LC n= 1, LG n= 1, LJ n= 1, LV n= 1 | 0 | - | - | - | - |
| PID156 | 25 | *LG n= 2* | 0 | - | - | - | - |
| PID157 | 32 | *LJ n= 1* | 0 | - | - | - | - |
| PID162 | 31 | *LC n= 4* | 0 | - | - | - | - |
| PID173 | 26 | *LC n= 6* | 0 | - | - | - | - |
| PID174 | 25 | *LJH n= 2* | 0 | - | - | - | - |
| PID179 | 18 | *LJ n= 1* | 0 | + | - | - | +/- |
| PID182 | 29 | *LC n= 3* | 0 | - | - | - | - |

PID: Patient identification number; ATCC: American Type Culture Collection; LC: *Lactobacillus crispatus*; LG: *L. gasseri*; LJ: *L. jensenii*; LJH: *L.*
*johnsonii*; LS: *L. salivarius*; LV: *L. vaginalis*; +/- Suggestive of candidiasis or budding yeast

### Lactic acid production and culture acidification

D- and L-lactate and pH were measured in culture supernatants of each isolate following overnight incubation under anaerobic conditions and the concentrations of lactic acid, a protective and immunoregulatory metabolite produced by lactobacilli (10, 25) were calculated. Lactic acid production and culture acidification were highly varied between species and strains (**Figure 1**). *L. crispatus* and *L. jensenii* isolates produced the most D-lactate (P=0.0033 and P=0.0152 for *L. crispatus* and *L. jensenii* versus *L. gasseri*, respectively; **Figure 1A**). A single *L. salivarius* strain and two *L. johnsonii* strains produced the most L-lactate, followed by *L. crispatus* (P=0.0047 for *L. crispatus* versus *L. jensenii*) (**Figure 1B**). *L. crispatus* also produced the most total lactate (P=0.0050 and P=0.0396 for *L. crispatus* versus *L. gasseri* and *L. vaginalis*, respectively; **Figure 1C**). As expected, total lactate was inversely correlated with culture pH (R = -0.8023; P<0.0001; **Figure 1E**). Similar differences were observed between species for lactic acid production, with *L. crispatus* producing the most D-lactic acid (P=0.0073 and P=0.0029 for *L. crispatus* versus *L. gasseri* and *L. vaginalis*, respectively; **Figure 1F**) and also producing the most L-lactic acid, other than the single *L. salivarius* strain (P=0.0011 and P=0.0125 for *L. crispatus* versus *L. jensenii* and *L. vaginalis*, respectively; **Figure 1G**).

**Figure 1:**
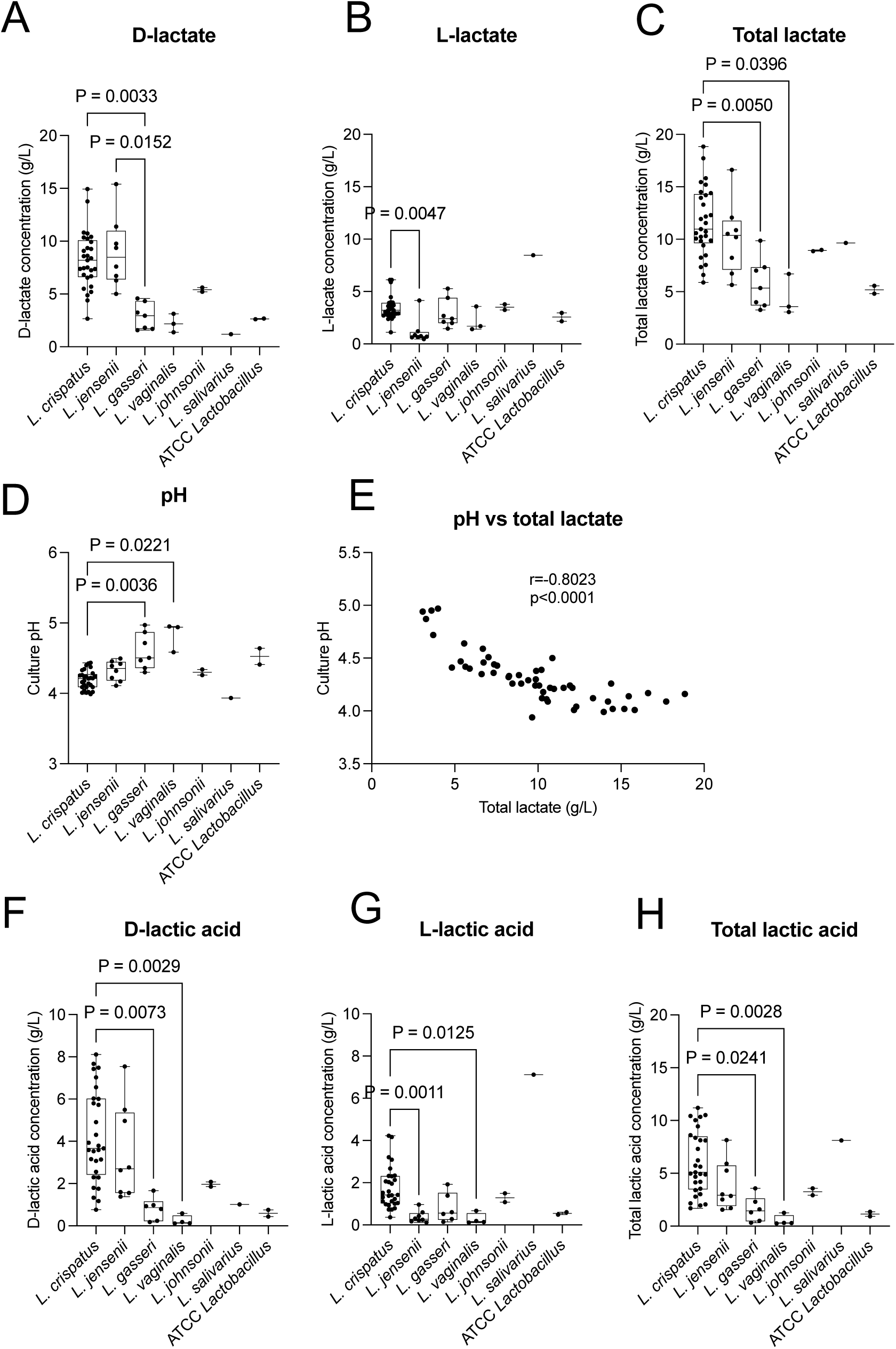
Lactate and lactic acid production and culture acidification by vaginal *Lactobacillus* isolates. D-lactate **(A)**, L-lactate **(B)**, total lactate **(C)** production, culture acidification **(D)**, the correlation between total lactate and pH **(E)**, D-lactic acid **(F)**, L-lactic acid **(G)** and total lactic acid production **(H)** are shown. Lactate and pH levels were measured in culture supernatants following 24 h incubation of standardised *Lactobacillus* isolates (*L. crispatus* (n=29), *L. jensenii* (n=8), *L. gasseri* (n=7), *L. vaginalis* (n=3), *L. johnsonii* (n=2), *L. salivarius* (n=1)) and two ATCC reference strains (*L. crispatus* ATCC® 33820^TM^ and *L vaginalis* ATCC® 49540^TM^) in de Man, Rogosa and Sharpe broth under anaerobic conditions. Lactic acid concentrations were calculated using the Henderson-Hasselbalch equation. Boxes represent the interquartile ranges, lines within boxes represent medians and whiskers represent minimum and maximum values. Each dot represents the average of two-three technical replicates. Kruskal-Wallis test and Dunn’s multiple comparisons test were used to compare characteristics between *Lactobacillus* species. Adjusted P-values <0.05 were considered statistically significant. ATCC: American Type Culture Collection.

### Inflammatory and anti-inflammatory cytokine responses to *Lactobacillus* isolates

The majority of inflammatory cytokine responses to lactobacilli were lower than those elicited by non-optimal *Gardnerella vaginalis* clinical isolates when co-cultured with vaginal epithelial cells, with significant differences noted between *G. vaginalis* strains and *L. crispatus* and *L. jensenii* (**Figure 2**). In contrast, lactobacilli elicited greater production of the anti-inflammatory cytokine IL-1RA compared to *G. vaginalis* at the 1 x 10^5^ colony forming units (CFU)/ml concentration, with significant differences noted between *G. vaginalis* and *L. crispatus* (P=0.0406) *and L. gasseri* (P=0.0122) (**Figure 2**). However, Some *Lactobacillus* isolates and the American Type Culture Collection (ATCC) reference strains elicited substantial inflammatory responses that were comparable to *G. vaginalis* (**Figure 3A**), albeit that 4.2 x 10^6^ CFU/ml lactobacilli were added compared to 1 x 10^6^ CFU/ml *G. vaginalis*, a lower concentration of *G. vaginalis* that was nonetheless sufficient to induce increased cell death. Only one *L. gasseri* isolate reduced cell viability to below 80% (**Figure 3B**). Overall, *L. crispatus* strains elicited the lowest inflammatory cytokine responses, relative to other lactobacilli. When cytokine responses and other variables (lactate, lactic acid, culture pH and VK2 cell viability) were compared between *Lactobacillus* isolates from STI positive versus negative women, no significant differences were found, although isolates from STI positive women tended to produce less L-lactate (p=0.0778; **Additional File 1, Supplementary Figure 1**).

**Figure 2:**
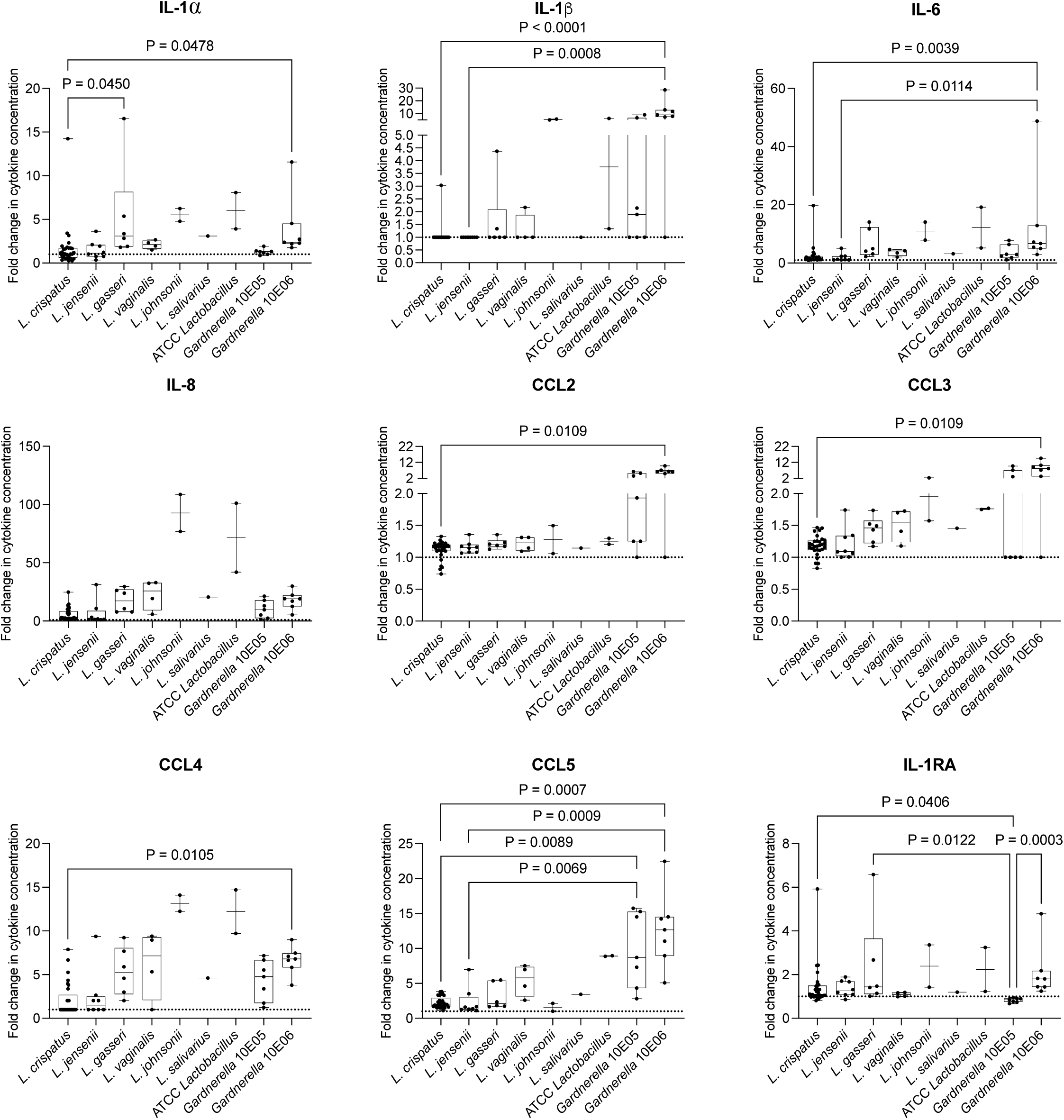
Fold changes in cytokine responses to *Lactobacillus* and *Gardnerella vaginalis* isolates and ATCC reference strains in an *in vitro* vaginal epithelial cell model. *Lactobacillus* and *G. vaginalis* isolates and ATCC reference strains (*L. crispatus* ATCC® 33820^TM^ and *L vaginalis* ATCC® 49540^TM^) were incubated with VK2 cells at 37°C, 5% CO_2_ for 24 h. *L. crispatus* (n=29), *L. jensenii* (n=8), *L. gasseri* (n=7), *L. vaginalis* (n=3), *L. johnsonii* (n=2), *L. salivarius* (n=1), *G. vaginalis* (n=7) were included. A total of 4.2 x 10^6^ colony forming units (CFU)/ml of each *Lactobacillus* strain was added, while two concentrations of *G. vaginalis* isolates were included as positive controls: 1 x 10^5^ CFU/ml and 1 x 10^6^ CFU/ml. After incubation, viability of the VK2 cells was assessed using the trypan blue assay and culture supernatants were collected for measurement of interleukin (IL)-6, IL-8, IL-1α, IL-1β, IL-1 receptor antagonist (RA), chemokine ligand (CCL)-2 (monocyte chemoattractant protein (MCP)-1), CCL3 (macrophage inflammatory protein (MIP)-1α), CCL4 (MIP-1β) and CCL5 (regulated on activation, normal T cell expressed and secreted (RANTES)) using Luminex. Fold changes in cytokine concentrations are shown for each isolate relative to untreated VK2 cells. Boxes represent the interquartile ranges, lines within boxes represent medians and whiskers represent minimum and maximum values. *Lactobacillus* isolates were evaluated in two independent experiments, as well as two Luminex technical replicates, and the averages for each strain shown as dots. Kruskal-Wallis test and Dunn’s multiple comparisons test were used to compare characteristics between *Lactobacillus* species. Adjusted P-values <0.05 were considered statistically significant. ATCC: American Type Culture Collection.

**Figure 3:**
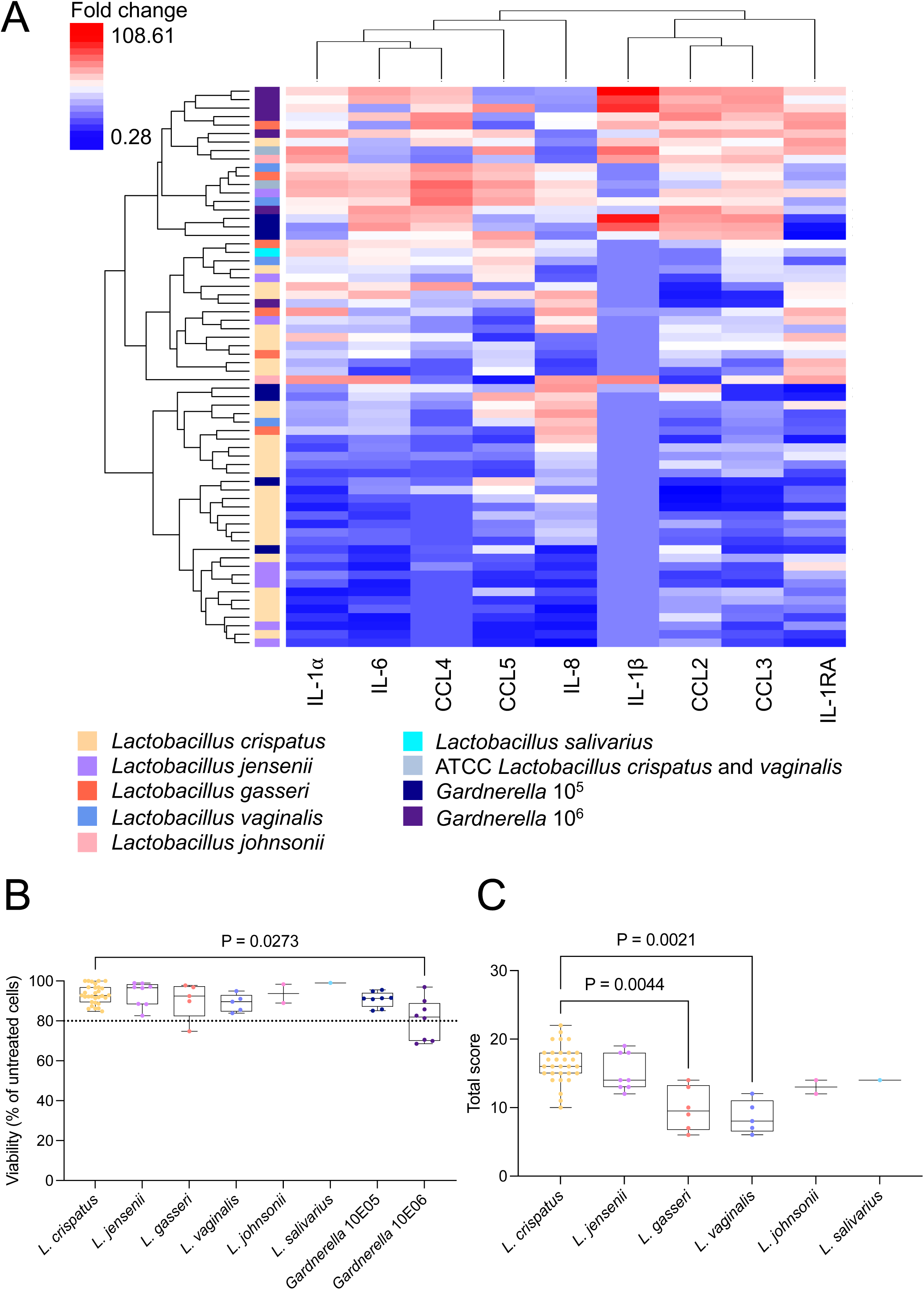
Cytokine responses to *Lactobacillus* and *Gardnerella vaginalis* isolates and ATCC reference strains, vaginal epithelial cell viability following co-culture and overall ranking of *Lactobacillus* species. (A) *Lactobacillus* and *G. vaginalis* isolates and ATCC reference strains (*L. crispatus* ATCC® 33820^TM^ and *L. vaginalis* ATCC® 49540^TM^) were incubated with VK2 cells at 37°C, 5% CO_2_ for 24 h. *L. crispatus* (n=29), *L. jensenii* (n=8), *L. gasseri* (n=7), *L. vaginalis* (n=3), *L. johnsonii* (n=2), *L. salivarius* (n=1), *G. vaginalis* (n=7) were included. After incubation, culture supernatants were collected for measurement of interleukin (IL)-6, IL-8, IL-1α, IL-1β, IL-1 receptor antagonist (RA), chemokine ligand (CCL)-2 (monocyte chemoattractant protein (MCP)-1), CCL3 (macrophage inflammatory protein (MIP)-1α), CCL4 (MIP-1β) and CCL5 (regulated on activation, normal T cell expressed and secreted (RANTES)) using Luminex. Fold changes in cytokine concentrations are shown for each isolate relative to untreated VK2 cells ranging from blue (low) through white to red (high). Two clustering dendrograms are shown in the figure. The dendrogram above the heat map illustrates degrees of relatedness between different cytokines measured. The dendrogram on the left hand side of the heat map indicates relationships between the expression profiles of the analysed cytokines in response to different bacteria. **(B)** The viability of the VK2 cells treated with lactobacilli or *G. vaginalis* was assessed using the trypan blue assay. **(C)** The overall biotherapeutic-relevant scores of each of the isolates are shown. Dots represent individual isolates; boxes represent the interquartile ranges, lines within boxes represent medians and whiskers represent minimum and maximum values. Kruskal-Wallis test and Dunn’s multiple comparisons test were used to compare characteristics between *Lactobacillus* species. Adjusted P-values <0.05 were considered statistically significant. ATCC: American Type Culture Collection.

### Overall ranking of *Lactobacillus* isolates according to biotherapeutic-relevant characteristics

All characteristics assessed (including D-lactate, L-lactate, culture acidification, inflammatory and anti-inflammatory cytokine production) were evaluated in combination to identify the best performing isolates (**Table 2**; **Figure 3C**). *L. crispatus* isolates performed better than the other species evaluated, and the majority of the top 10 isolates were *L. crispatus* strains (**Table 2**). Although isolates from women with STIs were included, there were no significant differences in the total ranking of isolates from women with and without STIs.

**Table 2.** Ranking of *Lactobacillus* isolates according to biotherapeutic-relevant characteristics.

| Isolate ID | Species | D-lactate | L-lactate | pH | Inflammation | IL-1RA | TOTAL |
| --- | --- | --- | --- | --- | --- | --- | --- |
| **PID182 3a** | ***Lactobacillus crispatus*** | 4 | 4 | 4 | 8 | 2 | 22 |
| **PID153 1a** | ***Lactobacillus crispatus*** | 4 | 3 | 4 | 8 | 2 | 21 |
| **PID150 3a** | ***Lactobacillus crispatus*** | 3 | 4 | 4 | 6 | 3 | 20 |
| PID150 5b | *Lactobacillus crispatus* | 4 | 4 | 3 | 6 | 3 | 20 |
| PID182 2b | *Lactobacillus crispatus* | 4 | 4 | 3 | 8 | 1 | 20 |
| PID157 1a | *Lactobacillus jensenii* | 4 | 1 | 4 | 8 | 2 | 19 |
| PID153 2a | *Lactobacillus jensenii* | 4 | 1 | 2 | 8 | 3 | 18 |
| **PID173 1a** | ***Lactobacillus crispatus*** | 4 | 2 | 4 | 6 | 2 | 18 |
| PID182 2a | *Lactobacillus crispatus* | 3 | 3 | 3 | 6 | 3 | 18 |
| **PID32 D1** | ***Lactobacillus crispatus*** | 3 | 2 | 3 | 8 | 2 | 18 |
| PID40 D2 | *Lactobacillus jensenii* | 3 | 4 | 1 | 6 | 4 | 18 |
| **PID41 D10** | ***Lactobacillus crispatus*** | 3 | 2 | 3 | 6 | 4 | 18 |
| PID43 5a | *Lactobacillus crispatus* | 4 | 4 | 4 | 2 | 4 | 18 |
| **PID162 5a** | ***Lactobacillus crispatus*** | 4 | 4 | 4 | 2 | 3 | 17 |
| PID173 1b | *Lactobacillus crispatus* | 2 | 3 | 2 | 8 | 2 | 17 |
| PID32 D9 | *Lactobacillus crispatus* | 3 | 2 | 2 | 8 | 2 | 17 |
| PID38 D6 | *Lactobacillus crispatus* | 2 | 4 | 4 | 6 | 1 | 17 |
| PID173 2a | *Lactobacillus crispatus* | 3 | 3 | 3 | 6 | 1 | 16 |
| PID32 D6 | *Lactobacillus crispatus* | 3 | 2 | 2 | 8 | 1 | 16 |
| PID38 D8 | *Lactobacillus crispatus* | 3 | 3 | 4 | 2 | 4 | 16 |
| PID41 D2 | *Lactobacillus crispatus* | 3 | 3 | 3 | 6 | 1 | 16 |
| PID43 2a | *Lactobacillus crispatus* | 2 | 2 | 3 | 8 | 1 | 16 |
| PID162 1a | *Lactobacillus crispatus* | 4 | 2 | 4 | 2 | 3 | 15 |
| PID162 3b | *Lactobacillus crispatus* | 3 | 3 | 3 | 2 | 4 | 15 |
| PID173 3b | *Lactobacillus crispatus* | 2 | 4 | 3 | 2 | 4 | 15 |
| **PID41 D4** | ***Lactobacillus crispatus*** | 2 | 1 | 2 | 6 | 4 | 15 |
| PID65 4b | *Lactobacillus crispatus* | 4 | 4 | 4 | 2 | 1 | 15 |
| PID150 4b | *Lactobacillus jensenii* | 4 | 1 | 3 | 2 | 4 | 14 |
| PID153 4b | *Lactobacillus gasseri* | 2 | 4 | 2 | 2 | 4 | 14 |
| PID173 2b | *Lactobacillus crispatus* | 3 | 2 | 3 | 2 | 4 | 14 |
| PID174 1a | *Lactobacillus johnsonii* | 2 | 4 | 2 | 2 | 4 | 14 |
| PID40 D1 | *Lactobacillus crispatus* | 2 | 3 | 1 | 6 | 2 | 14 |
| **PID40 D9** | ***Lactobacillus crispatus*** | 2 | 2 | 1 | 6 | 3 | 14 |
| PID41 D3 | *Lactobacillus jensenii* | 2 | 1 | 1 | 8 | 2 | 14 |
| PID42 D1 | *Lactobacillus salivarius* | 1 | 4 | 4 | 2 | 3 | 14 |
| PID150 2a | *Lactobacillus jensenii* | 4 | 1 | 3 | 2 | 3 | 13 |
| PID156 1a | *Lactobacillus gasseri* | 1 | 4 | 2 | 2 | 4 | 13 |
| PID43 3a | *Lactobacillus jensenii* | 2 | 1 | 1 | 8 | 1 | 13 |
| **PID162 2a** | ***Lactobacillus crispatus*** | 4 | 3 | 2 | 2 | 1 | 12 |
| PID174 3a | *Lactobacillus johnsonii* | 2 | 3 | 2 | 2 | 3 | 12 |
| PID179 1a | *Lactobacillus jensenii* | 3 | 1 | 2 | 2 | 4 | 12 |
| PID65 1b | *Lactobacillus vaginalis* | 1 | 3 | 1 | 6 | 1 | 12 |
| ATCC LC | *Lactobacillus crispatus* | 1 | 2 | 2 | 2 | 4 | 11 |
| ATCC LV | *Lactobacillus vaginalis* | 1 | 3 | 1 | 2 | 3 | 10 |
| **PID173 5a** | ***Lactobacillus crispatus*** | 1 | 3 | 2 | 2 | 2 | 10 |
| PID65 4a | *Lactobacillus gasseri* | 2 | 2 | 1 | 2 | 3 | 10 |
| PID156 3a | *Lactobacillus gasseri* | 1 | 2 | 1 | 2 | 3 | 9 |
| PID62 2a | *Lactobacillus vaginalis* | 1 | 2 | 1 | 2 | 2 | 8 |
| PID65 2b | *Lactobacillus gasseri* | 1 | 1 | 1 | 2 | 2 | 7 |
| PID79 2a | *Lactobacillus vaginalis* | 1 | 1 | 1 | 2 | 2 | 7 |
| PID153 3b | *Lactobacillus vaginalis* | 1 | 1 | 1 | 2 | 1 | 6 |
| PID38 D3 | *Lactobacillus gasseri* | 1 | 1 | 1 | 2 | 1 | 6 |

PID: Participant identity number; IL-1RA: Interleukin 1 receptor antagonist. Highlighted isolates were obtained from women who had sexually transmitted infections. Isolates in bold were analysed using whole genome sequencing.

### WGS analysis of *L. crispatus* isolates

The three most optimal *L. crispatus* isolates according to the characteristics assessed in this study were selected for WGS to evaluate additional live biotherapeutic relevant characteristics, including the presence of antimicrobial resistance (AMR) elements, putative prophages and bacteriocins. Seven additional *L. crispatus* isolates with varying performance were also included in this analysis for comparison. The average genome size of all ten *L. crispatus* isolates was 2.53 megabase pairs (Mbp; range: 2.45 Mbp to 2.58 Mbp) (**Additional File 2, Supplementary Table 1**). Phylogenetic analysis showed that the isolates clustered separately from the ATCC 33820 reference strain, two *Gallus gallus* (chicken) strains, as well as Human Microbiome Project isolates and a range of isolates from other geographical regions, including the US, Brazil, Netherlands and Italy (**Figure 4; Additional File 2, Supplementary Table 2**). Eight of the isolates formed a distinct clade that did not include any isolates from other geographical regions. Isolates from the same participants (PID162 2a and PID162 5a; PID173 1a and PID173 5a) were highly similar, with alignment percentages and average nucleotide identities of 99.14% and 100% for PID162 2a and PID162 5a, and 97.43% and 99,99% for PID173 1a and PID173 5a, respectively. This suggests that the same or similar strains were isolated multiple times and may thus be overrepresented in the microbiomes of these women (**Additional File 1, Supplementary Figure 2**).

**Figure 4:**
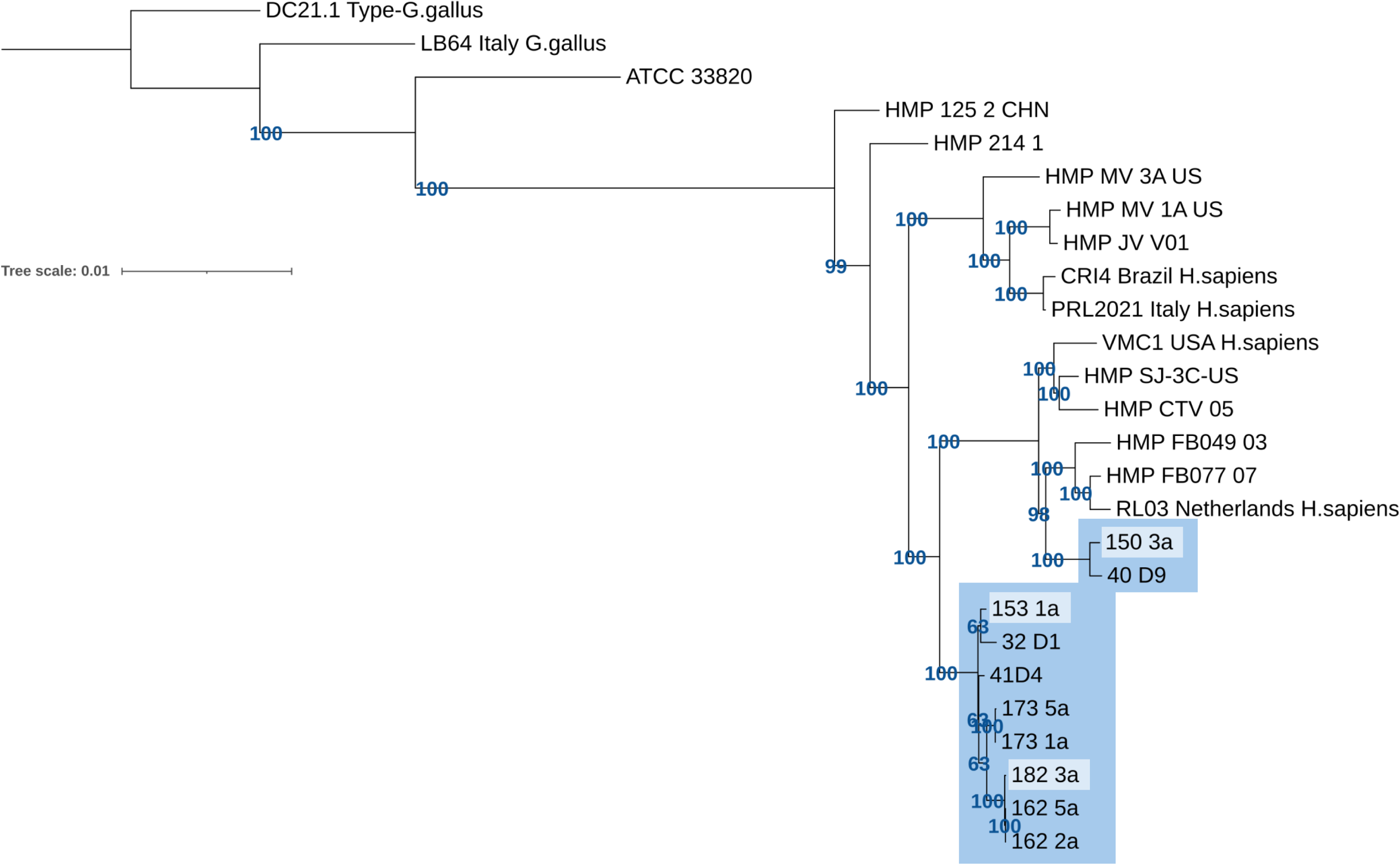
Estimated evolutionary relationships among South African *Lactobacillus crispatus* isolates, reference strains, and publicly available sequences for isolates from other geographical regions. Maximum likelihood phylogeny using the best-fit model (GTR+F+R2). Branches are scaled according to the number of nucleotide substitutions per site. Support for key nodes on the phylogeny are shown in the form of bootstrap support. All novel isolates are highlighted in blue, with the three most optimal isolates according to the assessed characteristics considered essential for optimal live biotherapeutic performance highlighted in light blue. Strains from other geographical regions, China (CHN), United States (US/USA), Brazil, Italy, and Netherlands, were included for comparison. including ATCC: American Type Culture Collection; HMP: Human Microbiome Project; *G. gallus*: *Gallus gallus*; *H. sapiens*: *Homo sapiens*.

No AMR genes or chromosomal mutations were detected in any *L. crispatus* assemblies using ResFinder (ver 4.6.0) and CARD (ver 3.3.0), regardless of isolate ranking. The presence of plasmids were also not detected in any *L. crispatus* assemblies via PlasmidFinder (ver 2.0.1). Although large differences were observed in lactic acid production between isolates, the L-lactate dehydrogenase and D-lactate dehydrogenase gene sequences were highly conserved, with 100% and 99.7% identity across all isolates, respectively. For D-lactate dehydrogenase, all isolates showed a single non-synonymous mutation compared to ATCC 33820, V331A. However, the functional importance of this mutation is unknown.

Bacteriocins were also of interest due to their integral role in inhibiting the growth of non-optimal bacteria in the vaginal microbiome (26). Overall, five putative bacteriocin-encoding genes were identified using BAGEL4 (**Additional File 2, Supplementary Table 3**). These included pediocin-like bacteriocin penocin A, the bacteriolysin enterolysin A, and two copies of helveticin J that were found across all isolates, as well as linear azol(in)e-containing peptides (LAPs) that was detected in 7/10 isolates.

The presence of prophage sequences within the assemblies was evaluated using PHASTEST (ver 3.0), checkV (ver. 1.0.3) and Pharokka (ver. 1.7.5). Intact putative prophages were identified across all isolates, however, for one isolate (PID40 D9), two prophages identified as intact using PHASTEST were classified as genome fragments using checkV (**Additional File 2, Supplementary Table 4**). While the putative prophages were most closely matched to *Listeria* phage B054 (all isolates), *Lactobacillus* phage Lc-Nu (two isolates), and *Lactobacillus* phage phiadh (seven isolates) in the PHASTEST database, the closest matches were identified as *Lactobacillus* phage vB_Lcr_AB1 and *Lactobacillus* phage vB_Lga_AB1 using both checkV and Pharokka (**Additional File 2, Supplementary Table 4**). Pairwise alignments showed varying sequence identity between the prophage sequences for the novel isolates compared to *Lactobacillus* phage vB_Lcr_AB1 (**Additional File 1, Supplementary Figure 3; Additional File 2, Supplementary Table 5**) and *Lactobacillus* phage vB_Lga_AB1 (**Additional File 1, Supplementary Figure 4; Additional File 2, Supplementary Table 6**), with some isolates displaying limited similarity. This suggests that the identified putative prophages may be only distantly related to prophages present in the reference databases included in this analysis. Only a single prophage sequence from 173 1a mapped to a smaller contig and may thus represent an induced prophage. No plasmids were identified in the data using PlasmidFinder. The number, type and completeness of the putative prophages identified were similar between the top three *L. crispatus* isolates and the remaining seven isolates.

## Discussion

This study characterised biotherapeutic-relevant properties of 50 novel vaginal *Lactobacillus* isolates from 18 young South African women. The primary outcome of this work is the expansion of the number of well-characterised optimal vaginal *Lactobacillus* isolates from African women to inform the development of effective live biotherapeutics for this population. This is critical because the vaginal microbiota differs by ethnicity and geographical region (20) and it is not known whether biotherapeutic strains from women of one region will effectively colonize women of another. Vaginal environmental factors are likely to vary between women of different ethnicities residing in different geographical regions, influenced by both genetic and external factors (e.g. hygiene practices, sexual enhancer insertion, diet, smoking, contraceptive choice) (27). These differences may result in strain-level variation of bacteria that have adapted to particular environments and this variation may in turn hinder growth and stable colonisation if utilised in different populations. In support of this, and similar to previous findings (23), it was found that the South African *L. crispatus* isolates subjected to WGS in this study were generally more closely related to one another than to reference strains and isolates from other geographical regions.

The key biotherapeutic-relevant characteristics that were evaluated, included D- and L-lactate and lactic acid production, culture acidification, pro- and anti-inflammatory cytokine production, and the presence of putative intact prophages, antimicrobial resistance elements, and bacteriocin genes. Lactic acid was evaluated as previous studies have shown that this metabolite has potent virucidal activity against different HIV subtypes at physiological concentrations (10, 28), with L-lactic acid more effective at inactivating HIV-1 than D-lactic acid at threshold concentrations and in a pH-dependent manner (10). On the other hand, D-lactic acid has been found to be more effective at inhibiting *C. trachomatis* infection of FGT epithelial cells than L-lactic acid (29). However, D-lactic acid and L-lactic acid have been observed to have similar anti-inflammatory and epithelial barrier strengthening activities on cervicovaginal epithelial cells *in vitro* (25, 30). Cytokine responses to the *Lactobacillus* isolates were also evaluated as women with inflammation in their genital tracts are at greater risk of HIV infection compared to those without inflammation (7). Each of the selected cytokines, except anti-inflammatory IL-1RA, has previously been associated with increased risk of HIV acquisition (7, 31). Varying inflammatory responses to the isolates observed in this study highlights the importance of assessing the inflammatory properties of all potential live biotherapeutics. Collectively, *L. crispatus* performed the best according to these characteristics, producing the most lactate, acidifying the culture media the most and inducing minimal inflammatory cytokine production. While *L. crispatus* isolates were the least inflammatory species overall, some isolates, as well as the ATCC reference lactobacilli, induced substantial inflammatory cytokine production. For this study, *G. vaginalis,* a key BV-associated species that causes inflammation both *in vivo* and *in vitro* and is associated with increased risk of HIV acquisition (2, 8, 32), was included as a positive control. Although the majority of *Lactobacillus* isolates elicited lower cytokine responses compared to *G. vaginalis* isolates, some were comparably inflammatory. The concentration of lactobacilli added (4.2 x 10^6^ CFU/ml) is within the range found *in vivo*, with previous studies showing that vaginal secretions contain between 10^3^ and 10^9^ CFU/ml of lactobacilli (33, 34). However, it should be noted that although >2 x 10^7^ CFU/ml of *G. vaginalis* can be found in women with BV (35). Higher *G. vaginalis* concentrations could not be utilised in this *in vitro* study as 10^6^ CFU/ml was sufficient to reduce cell viability. We have previously reported that lactobacilli from women with BV were more inflammatory than isolates from women without BV, suggesting significant variation in inflammatory properties within this genus (36).

Although the ten isolates included in the WGS analysis produced widely varying amounts of D- and L-lactic acid *in vitro*, D- and L-lactate dehydrogenase gene sequences were highly similar across all isolates. In addition to lactate dehydrogenase catalytic activity, other factors can influence the amount of lactic acid produced, including the expression of other enzymes that compete with lactate dehydrogenase for the utilization of pyruvate, efficiency of glucose import and lactate efflux and tolerance to low pH (37). Understanding the mechanisms underlying the varying production of lactic acid may inform screening strategies for optimal live biotherapeutics development.

Bacteriocins are antimicrobial peptides that are produced by *Lactobacillus* and other bacterial taxa and thought to play an important role in maintaining an optimal vaginal microbiome (26). Therefore, the presence of bacteriocin-encoding genes is considered a favourable property for live biotherapeutic candidates. All strains regardless of ranking were predicted to harbour bacteriocin-encoding genes including penocin A, enterolysin A, and two copies of helveticin J, while 7/10 isolates also had LAPs. Helveticin J and enterolysin A have been previously detected within vaginal *L. crispatus* isolates although their roles are not fully characterised (38). However, enterolysin A produced by *Enterococcus faecalis* has been shown to degrade bacterial cell walls of species of lactococci, enterocci as well as *Listeria innocua,* whilst helveticin J produced by *Lactobacillus helveticus* has displayed inhibitory effects against other lactobacilli (39, 40). Penocin A isolated from the genome of a strain of *Pediococcus pentosaceus* has been shown to inhibit the growth of *Listeria* and *Clostridium* spp. (41), although the role of penocin A within the vaginal microbiome should be further explored. The presence of putative LAPs, a class of bacteriocins that can be cytolytic or have potent narrow-spectrum antibacterial activity (42), in some strains, but not others, may offer competitive advantage against other non-optimal vaginal bacteria. Additionally, the inhibitory spectrum may vary between different strains and should be tested against non-optimal vaginal bacteria to determine their effectiveness in inhibiting pathogen growth(43).

AMR genes and/or chromosomal mutations as well as plasmids were not detected in the isolates sequenced in this study, an important safety consideration for live biotherapeutics development. *L. crispatus* isolates have however been found to be resistant to metronidazole, with varying susceptibility to clindamycin, and discordance between predicted and phenotypic resistance profiles has been demonstrated previously (44–46). Therefore, resistance mechanisms likely exist that were not detected in this analysis, suggesting that phenotypical evaluation of AMR is also important for live biotherapeutic candidate evaluation. Intact putative prophages were identified across all *L. crispatus* isolates using PHASTEST, with each isolate having more than one putative prophage sequence, although two prophages identified in one isolate were classified as incomplete using checkV. Prophage sequences have frequently been detected in vaginal *L. crispatus* isolates in previous studies, with an in silico analysis identifying prophages in 742 of 895 publicly available *Lactobacillaceae* genomes (47–50). However, the detection of multiple prophages in individual isolates was less common in this previous in silico analysis compared to the present study. The presence of prophages may be a safety concern for live biotherapeutic development, due to the possibility of enhanced virulence or transfer of genetic factors including AMR genes. However, very few of the prophages identified in the in silico analysis were found to harbour genes of clinical concern and inducible prophages have also been identified in commercial probiotics (51, 52). Regardless, further evaluation of the functionality of the putative prophages identified here is needed prior to further development of these *Lactobacillus* strains as live biotherapeutics.

Although VK2 cells closely resemble primary vaginal epithelial cells with respect to morphology and immunocytochemical characteristics (53), a limitation of this study was that the use of this transformed cell line does not fully recapitulate *in vivo* conditions. In the future, it would be informative to test these isolates in 3D models of vaginal epithelia, including primary epithelial cells from women of African ancestry. Another limitation is that half (9/18) of the women from whom the 50 isolates were obtained did not have repeat BV testing at multiple timepoints to confirm sustained optimal microbiota. While some isolates were collected from women who had STIs, we did not observe any significant differences in their final ranking compared to isolates from STI negative women. However, L-lactate production tended to be lower in isolates from STI positive versus negative women. This would be important to further investigate in future studies with larger sample sizes. Importantly, although these isolates were collected from a small group of women residing in a single region in South Africa, they will contribute towards the collection of geographically-diverse isolates.

In summary, significant species and strain-level differences in the biotherapeutic-relevant properties of the *Lactobacillus* isolates were observed. *L. crispatus* isolates produced the most lactic acid, caused the most significant reduction in culture pH and elicited the lowest inflammatory cytokine responses *in vitro*. The subset of *L. crispatus* isolates analysed using WGS were generally more similar to each other than to isolates from other geographical regions, supporting the need for live biotherapeutics to be tailored for the population of intended use. Putative bacteriocins and prophages were identified in all isolates analysed using WGS, while no AMR resistance elements were detected. These isolates will be further characterised towards the development of a potential live biotherapeutic to establish an optimal microbiome in the FGT, as an HIV prevention strategy for African women or alternatively to contribute to multi-strain biotherapeutics, including bacteria from different populations.

## Methods

### Study cohort

This study included cervicovaginal fluid samples and data collected from 25 women at the time of enrolment in the National Institutes of Health-funded Mucosal Injury from Sexual Contact (MISC) study in Cape Town, South Africa (R01AI128792). Inclusion criteria included being HIV negative, not pregnant, sexually active, not having taken antibiotics in the past month and no history of cervical disease. Women were asked to abstain from sexual activity and vaginal product use for two weeks prior to the enrolment visit. This study was approved by the University of Cape Town Human Research Ethics Committee (UCT HREC REF: 696/2017) and the Biomedical Research Ethics Committee (BREC) at the University of KwaZulu-Natal (BF504/17) in South Africa, the Seattle Children’s Research Institute Institutional Review Board (STUDY00000462) in the US and the Alfred Hospital Ethics Committee (75/21) in Australia and all methods were performed in accordance with the relevant guidelines and regulations. All women provided written informed consent. For women <18 years old, a parental consent waiver was obtained with approval from the ethics committees and the local Community Advisory Board affiliated with the Desmond Tutu Health Foundation (DTHF), through which recruitment was conducted. The South African Department of Health’s recommendations for circumstances in which it is ethically justifiable to waive parental consent were followed. A study inclusion criterion was that the girls and women were already sexually active and the study collected detailed information about sexual activity, sexual partners, gender-based violence, abuse and neglect. Obtaining parental consent would mean that adolescents at highest risk would possibly be excluded from the study. Given that enrolment in the study could offer these vulnerable girls some benefits, the requirement for parental consent could be considered unethical in these instances.

### BV and STI diagnostics

Lateral vaginal wall swabs were collected for Nugent scoring by Gram staining of vaginal smears and STI testing (*Chlamydia trachomatis, Trichomonas vaginalis* and *Neisseria gonorrhoeae*) using Primerdesign^TM^ genesig® kits at Bio Analytical Research Corporation South Africa (BARC SA).

### Bacterial isolation and culture

For *Lactobacillus* isolation, cervicovaginal secretions were collected from women who had optimal microbiota (Nugent BV negative; Nugent score 0-3) using menstrual cups (Softcup®, US). Menstrual cup samples were placed at 4°C and transferred to the laboratory for processing within 6 h of collection. Samples were diluted 1:5 in phosphate buffered saline (PBS) and stored at -80°C in 26% glycerol in PBS. Cervicovaginal samples (n=25) were streaked on pre-reduced de Man, Rogosa and Sharpe agar (MRS; Merck Pty Ltd) and incubated anaerobically in a Don Whitley A35 workstation at 37°C for 48-72 h. Single colonies were re-streaked thrice on pre-reduced MRS agar and incubated anaerobically at 37°C for 48-72 h to ensure purity. Seven *G. vaginalis* strains were also isolated from menstrual cup samples from different women within the same cohort, including three BV negative and four BV positive women. For this, Columbia Horse Agar with 5% Horse Blood (CBA) (Edwards Group, Australia) and Brucella Blood Agar with Hemin and Vitamin K1 included (University of Melbourne, Australia) were used as above.

### Gram staining and 16S rRNA gene sequencing

For Gram staining, a smear of a bacterial colony of each isolate was applied to a glass slide and heat fixed. For 16S rRNA gene sequencing, a loopful of a bacterial colony of each isolate was emulsified in PrepMan™ Ultra Sample Preparation Reagent (Thermofisher), vortexed and incubated at 100°C for 10 min in a heating block. Isolates were then identified using 16S rRNA gene Sanger sequencing of the V1-V4 region at the Australian Genome Research Facility, Melbourne. Sequences were BLAST searched against the Greengenes database (https://greengenes.secondgenome.com).

### *Lactobacillus* spp. culture and standardisation

Of the 181 *Lactobacillus* spp. isolated, 50 were chosen based on unique morphological characteristics and microbial 16S rRNA gene identification. They included *L. crispatus*, *L. gasseri, L. vaginalis, L. jensenii, L. salivarius and L. johnsonii.* Two reference strains were included as controls: *L. crispatus* ATCC® 33820^TM^ and *L vaginalis* ATCC® 49540^TM^. As each experiment required the addition of a consistent number of bacteria, the numbers of viable bacteria present in cultures with an optical density (OD) at 600 nm of 0.1±0.01 were determined. Prior to each experiment, MRS broth was inoculated with *Lactobacillus* isolates and incubated for 24 h at 37°C under anaerobic conditions in a Don Whitley A35 workstation. Cultures were then adjusted to 4.2 x 10^6^ colony forming units (CFU)/mL in MRS broth based on OD_600_ readings as previously described (27, 31, 32), then incubated again for 24 h at 37°C anaerobically. For cytokine assays, the *G. vaginalis* strains were cultured in Brain Heart Infusion (BHI) broth (Edwards Group, Australia) under anaerobic conditions for 48-72h at 37°C anaerobically.

### Assessment of D-lactate, L-lactate, pH and lactic acid production

Following incubation of standardised bacterial cultures for 24 h, cultures were centrifuged at 6000xg for 5 minutes and supernatants were filtered using 0.22 μm Costar® Spin-X® centrifuge tubes prior to measurement of both lactate isomers using the D- and L-lactic acid kit (R-biopharm, Darmstadt, Germany) (28). pH was measured in culture supernatants using an Aqua pH meter (TPS) and total lactic acid was calculated using the Henderson-Hasselbach formula (28). For each isolate, two-three dilutions (1:10, 1:20 and 1:40) were evaluated across one independent assay and the averages of final calculated concentrations determined for all measurements within range. The intraassay reproducibility of the assay was high (R=0.99, p<0.0001 and R=0.85, p<0.0001 for D- and L-lactate, respectively), while lactate concentrations correlated for separate cultures of the same isolates (n=6) conducted by different technicians one year apart (R=0.72, p=0.05 and R=0.93, p=0.004, one-tailed, for D- and L-lactate, respectively).

### Inflammatory and immunoregulatory cytokine production

Vaginal epithelial cells (VK2/E6E7 ATCC® CRL-2616^™^) were maintained in complete keratinocyte serum free media (KSFM) supplemented with 0.4 mM calcium chloride, 0.05 mg/ml of bovine pituitary extract, 0.1 ng/ml human recombinant epithelial growth factor and 50 IU/ml penicillin and 50 µg/ml streptomycin (Sigma-Aldrich). The VK2 cells were seeded into 24-well tissue culture plates, incubated at 37°C in the presence of 5% CO_2_ and grown to confluency. Prior to bacterial addition, cells were washed 3x with antibiotic-free KSFM.

As previously described, lactobacilli were adjusted to 4.2 x 10^6^ CFU/ml in antibiotic-free KSFM and independently added to VK2-cell monolayers before incubating for 24 h at 37°C in the presence of 5% CO_2_ (36, 54). *G. vaginalis* isolates were adjusted to a similar concentration as the lactobacilli (1 x 10^6^ CFU/ml), but a 10-fold lower dose (1 x 10^5^ CFU/ml) was also included as some *G. vaginalis* strains are more virulent than others, as previously described (32). Co-culture supernatants were collected from each well for cytokine measurement after 24h of incubation. IL-6, IL-8, IL-1α, IL-1β, IL-1 receptor antagonist (RA), CCL2 (monocyte chemoattractant protein (MCP)-1), CCL3 (macrophage inflammatory protein (MIP)-1α), CCL4 (MIP-1β) and CCL5 (regulated on activation, normal T cell expressed and secreted (RANTES)) concentrations were measured using a Magnetic Luminex Screening Assay kit (R&D Systems). Data were collected using a Bio-Rad Bio-Plex® 200 system and a 5-parameter logistic regression was used to calculate cytokine concentrations from the standard curves using BIO-plex manager software (version 6.1; Bio-Rad Laboratories Inc®). Cytokine concentrations below the detectable limit were assigned the value of half the lowest recorded concentration of that cytokine. *Lactobacillus* isolates were evaluated in two independent experiments, as well as two Luminex technical replicates and the averages of three measurement for each isolate were analysed. VK2 cell viability following bacterial stimulation in one independent experiment was confirmed using the Trypan blue exclusion assay.

### Statistical analysis

Data were analysed using GraphPad Prism version 9 and R Studio version 1.2.1335. Distribution of variables was assessed by Shapiro-Wilk test. Kruskal-Wallis test with Dunn’s multiple comparisons test were used for unmatched comparisons and Spearman Rank test was used to test for correlations for non-parametric data. Unsupervised hierarchical clustering was used to evaluate overall cytokine production by VK2 cells in response to lactobacilli and *G. vaginalis*. Mann Whitney U test was used to evaluate differences between isolates from women with or without STIs. A false discovery rate (FDR) step-down procedure was used to adjust p-values for multiple comparisons for Spearman Rank and Mann Whitney U tests, with adjusted p-values <0.05 considered statistically significant.

### Isolate ranking

Isolates were given a maximum score of four per characteristic, including D-lactate, L-lactate and anti-inflammatory cytokine (IL-1RA) production. Relative scores per category were assigned as follows: <25^th^ percentile (score=1); 25^th^-50^th^ percentile (score=2); 50^th^-75^th^ percentile (score=3); ≥75^th^ (score=4). Lower culture pH levels were considered advantageous, so isolates were scored as follows: <25^th^ percentile (score=4); 25^th^-50^th^ percentile (score=3); 50^th^-75^th^ percentile (score=2); ≥75^th^ (score=1). For inflammatory cytokine induction, lower levels were also considered advantageous, and, since inflammatory profiles were considered critical, scores were doubled: <25^th^ percentile (score=8); 25^th^-50^th^ percentile (score=6); 50^th^-75^th^ percentile (score=4); ≥75^th^ (score=2).

### Whole genome sequencing analyses of isolates

Frozen bacterial isolates were inoculated in 4ml New York City (III) broth and incubated for 48-72 hours under anaerobic conditions. Bacterial cultures were then centrifuged at 2500 rpm for 5 minutes, followed by the collection of the bacterial cell pellets which were then frozen at -80°C. Bacterial cell pellets were then delivered to the Australian Genome Research Facility (AGRF, Melbourne, Victoria, Australia) for DNA extraction, library preparation and PacBio sequencing (Pacific Biosciences of California, Menlo Park, CA, USA). DNA extraction was performed via the Qiagen DNeasy PowerSoil Pro Kit (Qiagen, Hilden, Germany). Sequencing libraries were constructed via the SMRTbell® prep kit 3.0 (Pacific BioSciences, Menlo Park, California, USA) and long reads were sequenced on a PacBio Revio™ Platform with a single-molecule real-time (SMRT) 25M cell. HiFi reads were then assembled via the SMRT Link software (ver 13.1) to generate bacterial whole genome sequences according to the PacBio Microbial Whole Genome Sequencing Workflow. The reads were also assembled using Hybracter (55) for comparison, with both approaches yielding similar results.

Rapid and standardised annotation of whole genome assemblies was conducted using Bakta (ver 1.9.4) (56). Following annotation, assemblies were screened for genes and/or chromosomal mutations conferring for antimicrobial resistance using ResFinder (ver 4.6.0) (57) and the Comprehensive Antibiotic Resistance Database (CARD) (ver 3.3.0) (58) under the ‘Perfect and Strict hits only’ as well as the ‘Exclude nudge ≥95% identity Loose hits to

Strict’ settings. The presence of plasmids were screened for using PlasmidFinder (ver 2.0.1)(59, 60). The identification of prophage sequences within the isolates were determined via PHASTEST (ver 3.0)(61) under the ‘Deep’ setting. Prophage sequences obtained from PHASTEST were further examined through CheckV (ver. 1.0.3) (62) and Pharokka (ver 1.7.5) (63). Alignment and visualisation of homologous gene clusters between sequences obtained from CheckV and their top CheckV phage hit was conducted via clinker (64). Specific genes of interest including L-lactate dehydrogenase (GenBank accession number: QWW28174) and D-lactate dehydrogenase (GenBank accession number: KRK33509) were detected via BLASTN (ver 2.15.0+) (65). Gene clusters encoding putative bacteriocins were detected via the online web server BAGEL4 (66).

For phylogenetic analysis, genome sequences for *L. crispatus* for comparison were obtained from the National Center for Biotechnology Information (NCBI) (https://ftp.ncbi.nlm.nih.gov/genomes/HUMAN_MICROBIOM/Bacteria/ and

https://www.ncbi.nlm.nih.gov/datasets/genome/) in April 2025 (50, 67–69). *L. crispatus* genomes were aligned with the reference *L. crispatus* ATCC 33820 strain (GenBank: NZ_CP072197.1) using the Whole Genome Alignment tool within CLC Genomics (Qiagen, Hilden, Germany). Following alignment, regions of ambiguous and uncertain alignment were removed using Gblocks (70). Phylogenetic relationships were estimated using the maximum likelihood method, using the best-fit model (GTR+F+R2), with 1,000 ultrafast bootstrap replicates optimised for branch length with Nearest Neighbour Interchange using IQTree2 (71). The phylogenetic tree was visualised using iTOL (72).

## Availability of data and material

The datasets used and/or analysed during this study are available from the corresponding author on reasonable request. Raw whole genome sequence data and assemblies will be deposited at the European Bioinformatics Institute (http://www.ebi.ac.uk/) upon acceptance of the manuscript.

## Acknowledgments

The authors would like to acknowledge the women enrolled in the MISC cohort, as well as the clinical study staff. This work was supported by the National Institutes of Health (1R01AI128792), Carnegie Corporation of New York, Australian Centre for HIV and Hepatitis Virology Research and Joe White Bequest. LM was supported by the National Research Foundation (NRF) of South Africa and the Australian National Health and Medical Research Council. The funders had no role in study design, data collection and interpretation, or the decision to submit the work for publication. We gratefully acknowledge the contribution of the Victorian Operational Infrastructure Support Program, received by the Burnet Institute.

## Authors’ contributions

JW and ASAH conducted the laboratory work, analysed the data and contributed to manuscript preparation. CJ, MTM, DT, AGA, NR, RH, BM conducted some of the laboratory work and contributed to manuscript preparation. CML, HH, PG, LGB managed the clinical sites for the MISC study, collected some of the clinical data and contributed to manuscript preparation. JH, BK, AS, MZ contributed to data analysis and manuscript preparation. ACH, JSP, HJ, GT supervised the data collection and analysis and contributed to manuscript preparation. LM conceptualized the study, supervised the data collection, analysed some of the data and wrote the manuscript. All authors contributed to manuscript preparation.

## Ethics approval and consent to participate

This study was approved by the University of Cape Town Human Research Ethics Committee (UCT HREC REF: 696/2017) and the Biomedical Research Ethics Committee (BREC) at the University of KwaZulu-Natal (BF504/17) in South Africa, the Seattle Children’s Research Institute Institutional Review Board (STUDY00000462) in the US and the Alfred Hospital Ethics Committee (75/21) in Australia and all women provided written informed consent.

For women <18 years old, a parental consent waiver was obtained with approval from the ethics committees and the local Community Advisory Board affiliated with the Desmond Tutu Health Foundation (DTHF), through which recruitment was conducted. The South African Department of Health’s recommendations for circumstances in which it is ethically justifiable to waive parental consent were followed. A study inclusion criterion was that the girls and women were already sexually active and the study collected detailed information about sexual activity, sexual partners, gender-based violence, abuse and neglect. Obtaining parental consent would mean that adolescents at highest risk would possibly be excluded from the study, and that enrolment in the study could offer these vulnerable girls some benefits, and therefore, the requirement for parental consent could be considered unethical in these instances.

